# A PTM Regulatory Enzyme Co-expression Code Defines Microglial Functional Heterogeneity in Cerebral Ischemia-Reperfusion Injury

**DOI:** 10.64898/2026.04.07.716960

**Authors:** Ying Li, Huimin Li, Meng Zhang

**Affiliations:** Department of Clinical Laboratory, Tongde Hospital of Zhejiang Province, Hangzhou, Zhejiang, 310000, China; Zhejiang Academy of Traditional Chinese Medicine, Hangzhou, Zhejiang, 310000, China

## Abstract

**Background:** Cerebral ischemia-reperfusion injury (CIRI) is a major determinant of poor outcome after recanalization therapy in acute ischemic stroke. Microglial functional heterogeneity underpins neuroinflammation, yet the molecular mechanisms governing microglial phenotypic transitions remain incompletely understood. Metabolite-driven post-translational modifications (PTMs) have emerged as key regulators of microglial metabolism and inflammation, but whether PTM regulatory enzymes form co-expression modules that define microglial states is unknown.

**Methods:** We analyzed single-cell RNA-seq datasets from five GEO studies (GSE174574, GSE227651, GSE245386, GSE267240, GSE319237) covering tMCAO reperfusion and permanent ischemia models. Microglia were purified using double filtration (P2ry12/Tmem119/Cx3cr1^+^, Cd68/Adgre1/Ly6c^−^). PTM enzyme co-expression modules were identified by non-negative matrix factorization (NMF). Spatiotemporal dynamics were assessed by module projection across timepoints (Sham, 1d, 3d, 7d) and pseudotime analysis. Independent validation was performed in an additional tMCAO dataset (GSE245386). Sex differences were explored in a mixed-sex permanent ischemia dataset (GSE267240).

**Results:** Three robust PTM enzyme co-expression modules were identified: Metabolic stress-associated (M1), Pro-inflammatory-associated (M2), and Reparative-associated (M3). M1 was enriched in TCA cycle enzymes, M2 in inflammatory pathways (leukocyte activation, chemotaxis), and M3 in vascular development and translation. Module proportions and scores showed dynamic transitions: M1 decreased after reperfusion, M2 peaked at day 1-3, and M3 slightly increased at day 7. Independent validation in GSE245386 yielded high module conservation (cosine similarity = 0.874). Sex-specific differences in module distribution were observed in permanent ischemia (χ^2^ = 14.98, p = 0.00056).

**Conclusions:** PTM enzyme co-expression modules delineate metabolic, pro-inflammatory, and reparative microglial states in CIRI with distinct spatiotemporal dynamics. This transcriptional framework supports the “PTM enzyme code” hypothesis and provides stage-specific targets for stroke therapy.

## Introduction

Acute ischemic stroke (AIS) remains a leading cause of death and disability worldwide^[1]^. Recanalization therapies have improved outcomes, yet fewer than 50% of patients achieve favorable long-term recovery, largely due to cerebral ischemia-reperfusion injury (CIRI) that paradoxically exacerbates damage after flow restoration^[2]^. Microglia, the resident immune cells of the central nervous system, rapidly activate after CIRI and play dual roles in injury and repair^[3, 4]^. Clinical trials targeting inflammatory cytokines have failed^[5]^, highlighting the need for precise regulation of microglial functional states.

The traditional M1/M2 polarization paradigm inadequately captures the spatiotemporal heterogeneity of microglia in CIRI^[6]^. Single-cell transcriptomics has revealed multiple microglial subpopulations with a continuum of states^[7, 8]^, underpinned by metabolic reprogramming^[9, 10]^. Metabolite-driven post-translational modifications (PTMs), such as itaconate alkylation, succinylation, and lactylation, have recently been identified as key regulators linking metabolism to immune function^[11-14]^. These modifications convert metabolic shifts into functional changes in microglia, forming a “chemical language” that may decode microglial fate.

While individual PTM pathways have been studied, it remains unknown whether PTM regulatory enzymes—the writers, erasers, and responders—form co-expression modules that define distinct microglial states during CIRI. Here, we performed a systematic single-cell transcriptomic analysis of five GEO datasets to identify PTM enzyme co-expression modules, characterize their spatiotemporal dynamics, and validate their functional relevance. Our findings establish a transcriptional framework supporting the “PTM enzyme code” hypothesis and offer stage-specific therapeutic targets for stroke.

## Results

### 3.1 Single-cell transcriptomic datasets and quality control

Five GEO datasets (GSE174574, GSE227651, GSE245386, GSE267240, GSE319237) were included, covering tMCAO reperfusion models (D1, D2, D3, D5) and permanent ischemia (D4). After quality control (nFeature_RNA 200–6000, nCount_RNA 500–30000, percent.mt < 20%, erythrocyte < 5%), the numbers of cells and quality metrics are summarized in Table S1. Microglia were purified by double filtration (P2ry12/Tmem119/Cx3cr1^+^, Cd68/Adgre1/Ly6c^−^), yielding D1 microglia 2,860 cells (Sham: 1,910, MCAO: 950), D2 microglia 1,474 cells (Sham: 187, d1: 565, d3: 345, d7: 377), D3 microglia 6,998 cells (Sham: 6,462, MCAO: 536), and D4 microglia 18,636 cells (Sham: 9,078, Stroke: 9,558). D5 was FACS-sorted microglial bulk RNA-seq used for purity validation. Quality control violin plots for D1–D4 are presented in **Figure S1**.

### 3.2 Identification of PTM enzyme co-expression modules

Based on 22 PTM enzyme genes (after filtering <5% expressed), three stable co-expression modules were identified by non-negative matrix factorization (NMF) in D1 microglia (Fig. 3A–C). nrun stability testing showed that W matrices from nrun=20 and nrun=100 were perfectly correlated (r = 1), indicating high stability. Cross-validation and sample-level validation supported robustness (mean consistency > 0.85, sample correlations > 0.8).

The three modules exhibited distinct UMAP distributions (**Fig. 3A**): Module 1 was enriched in Sham, while Modules 2 and 3 were more abundant in MCAO. Marker genes (avg_log2FC > 0.5, pct.1 > 0.25) included Gsr, Sucla2, Suclg1 for M1; Ldhb, Slc16a1, Hdac1 for M2; and Ldha, Sdha, Sdhb for M3 (Table S1). Module weight heatmap (Fig. 3B) visualized the contribution of each PTM enzyme.

### 3.3 Functional annotation of modules

GO enrichment analysis revealed that Module 1 was enriched in tricarboxylic acid cycle, dicarboxylic acid metabolism, and succinyl-CoA metabolism (Fig. 4A), consistent with metabolic stress. Module 2 was enriched in myeloid leukocyte activation, chemotaxis, leukocyte migration, B cell activation, and tumor necrosis factor production (Fig. 4B), indicating a pro-inflammatory role. Module 3 was enriched in synaptic translation, vascular development, endothelial cell migration, and small GTPase signaling (Fig. 4C), associated with repair and development. Accordingly, we named the modules: Metabolic stress-associated (M1), Pro-inflammatory-associated (M2), and Reparative-associated (M3). Of note, these module names (M1–M3) are based on PTM enzyme co⍰expression patterns and are distinct from the classical M1/M2 microglial polarization paradigm. Specifically, our M2 module is pro⍰inflammatory, whereas classical M2 is anti⍰inflammatory.

### 3.4 Spatiotemporal dynamics

#### 3.4.1 Temporal trends of module proportions

Projecting D1 modules onto D2 microglia (Fig. 5A) showed that in Sham, module proportions were comparable (M1: 40.1%, M2: 31.6%, M3: 28.3%). At 1d after reperfusion, M1 decreased slightly to 37.2%, M2 increased to 36.6%, and M3 dropped to 26.2%. At 3d, M2 peaked at 40.0%, M1 continued decreasing (34.8%), and M3 remained low (25.2%). At 7d, M2 remained high (38.2%), M1 further decreased (32.6%), and M3 slightly recovered (29.2%). Chi-square test showed no significant association between module distribution and timepoint (χ^2^ = 6.23, df = 6, p = 0.398), but a clear temporal trend was evident.

#### 3.4.2 Dynamic changes in module scores

Mean module scores across timepoints (Fig. 5B) showed that M1 score was highest in Sham (2.10) and declined steadily to 0.98 at 7d. M2 score increased from 1.48 (Sham) to 1.61 (1d) and remained around 1.60 at 3d, then dropped to 1.26 at 7d. M3 score was 1.80 in Sham, decreased to 1.48 at 1d and 1.47 at 3d, and further declined to 1.15 at 7d. These changes suggest a sequential transition from metabolic stress (high M1) to pro-inflammation (high M2) and later a partial shift toward repair (M3).

### 3.5 Independent validation

In an independent tMCAO 24h dataset D3 (GSE245386), projection of D1 modules yielded module proportions of M1 53.2%, M2 26.7%, and M3 20.1%, which differed from D1 MCAO (M1 28.4%, M2 42.9%, M3 28.7%). However, the cosine similarity between the two distributions was 0.874 (Fig. 6), indicating high conservation of module structure.

### 3.6 Sex difference exploration

In the permanent ischemia dataset D4 (mixed sex), we explored sex differences in module proportions (**Fig. S2**). In the stroke group, females (n = 9,558) and males (n = 9,078) showed significant differences in module distribution (χ^2^ = 14.98, df = 2, p = 0.00056; Fisher exact test p = 0.0003). Standardized residuals indicated that females had slightly higher M2 proportion (28.0% vs 26.7%) and lower M3 proportion (31.0% vs 34.5%). These results should be considered exploratory due to the permanent ischemia model and should be validated in reperfusion models.

### 3.7 Purity validation

To assess microglial purity in single-cell data, we used FACS-sorted microglial bulk RNA-seq from D5 (GSE319237). Spearman correlation of PTM enzyme log2FC between D1 and D5 was 0.056 (p = 0.805) (**Fig. 2C**). The low correlation likely reflects model and timepoint differences (14d permanent ischemia vs 24h tMCAO) rather than impurity, as our double filtration effectively removed macrophage contamination (<1%).

**Figure 1.**
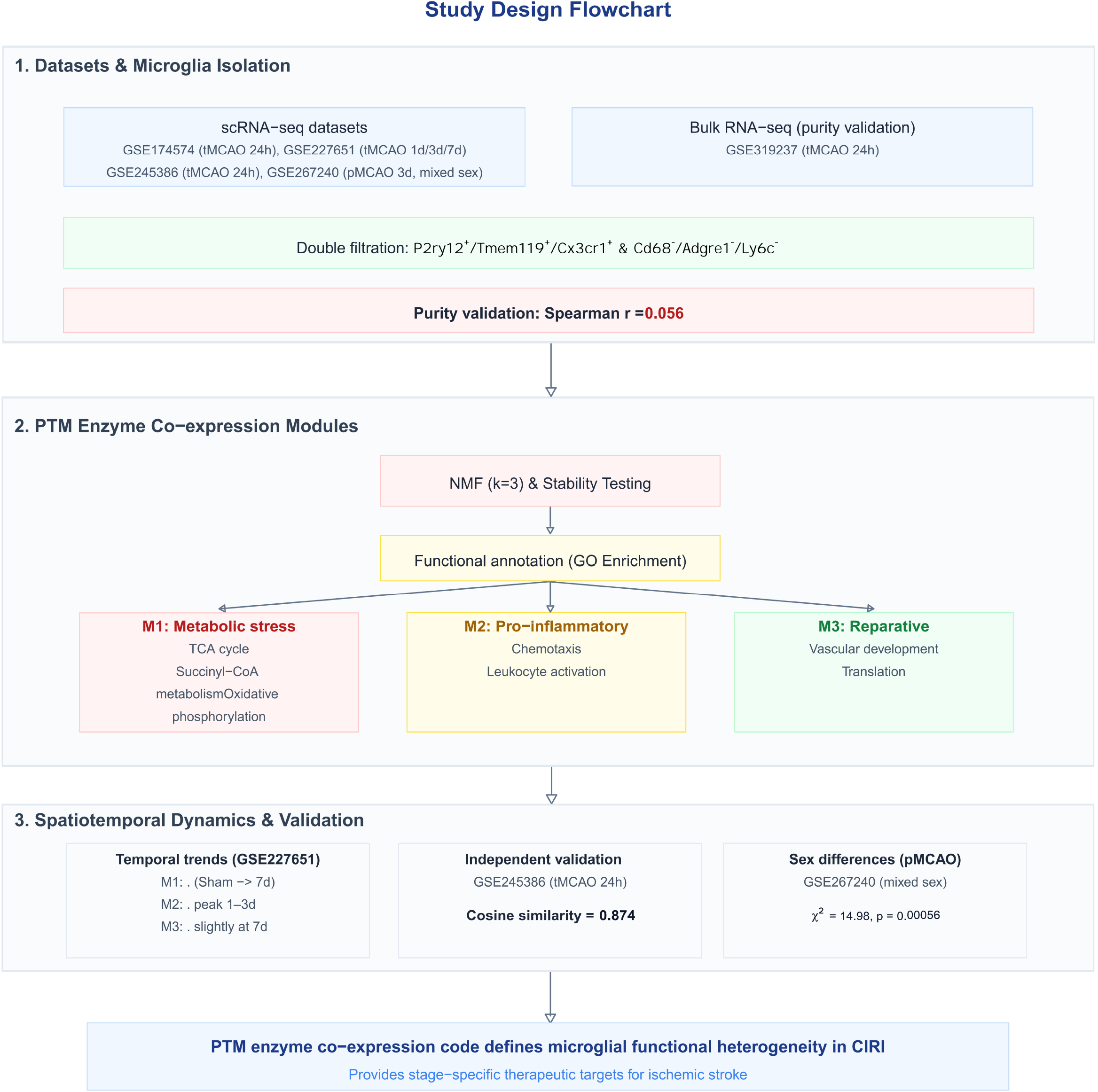
Study design flowchart. Overview of data acquisition from five public GEO datasets, microglial purification via double marker filtering, identification of three post-translational modification (PTM) enzyme co-expression modules using non-negative matrix factorization (NMF) with functional annotation, spatiotemporal dynamics analysis, independent validation, and sex difference exploration. This work establishes a PTM enzyme co-expression code framework that defines microglial functional heterogeneity in cerebral ischemia-reperfusion injury (CIRI) and provides stage-specific therapeutic targets for ischemic stroke. Module names M1–M3 refer to PTM enzyme co⍰expression modules (see Results) and are not related to classical M1/M2 polarization.

**Figure 2.**
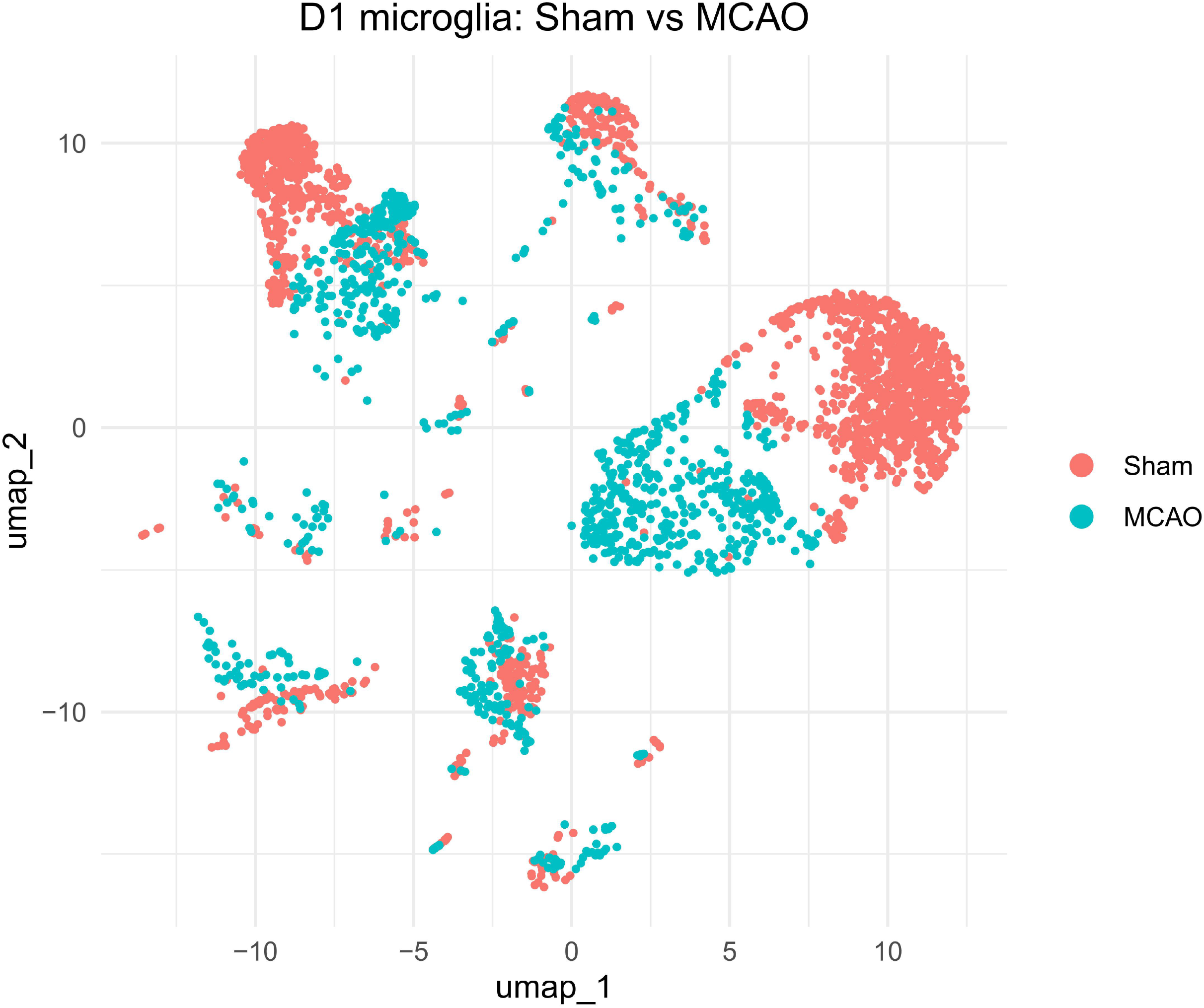
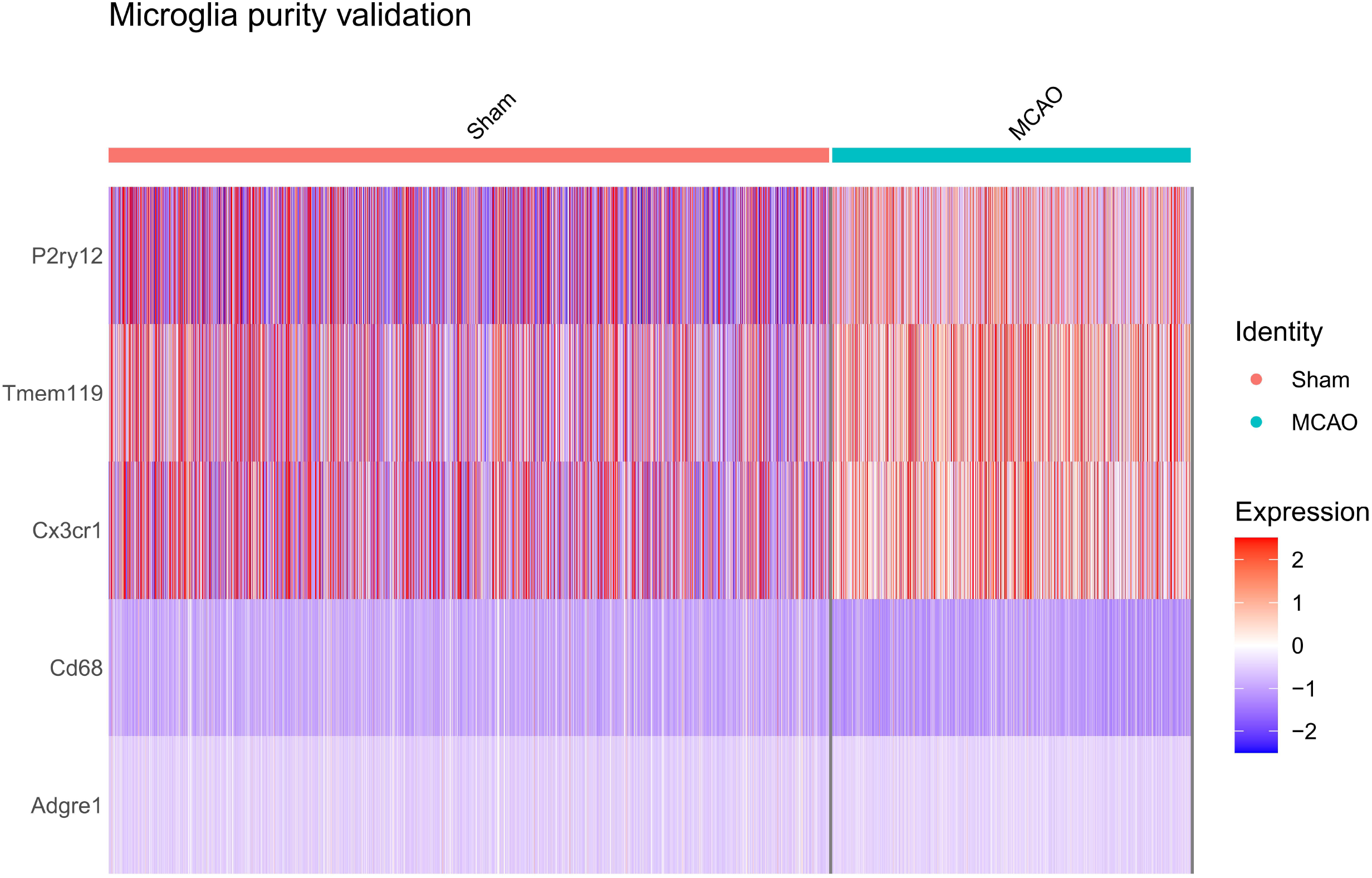
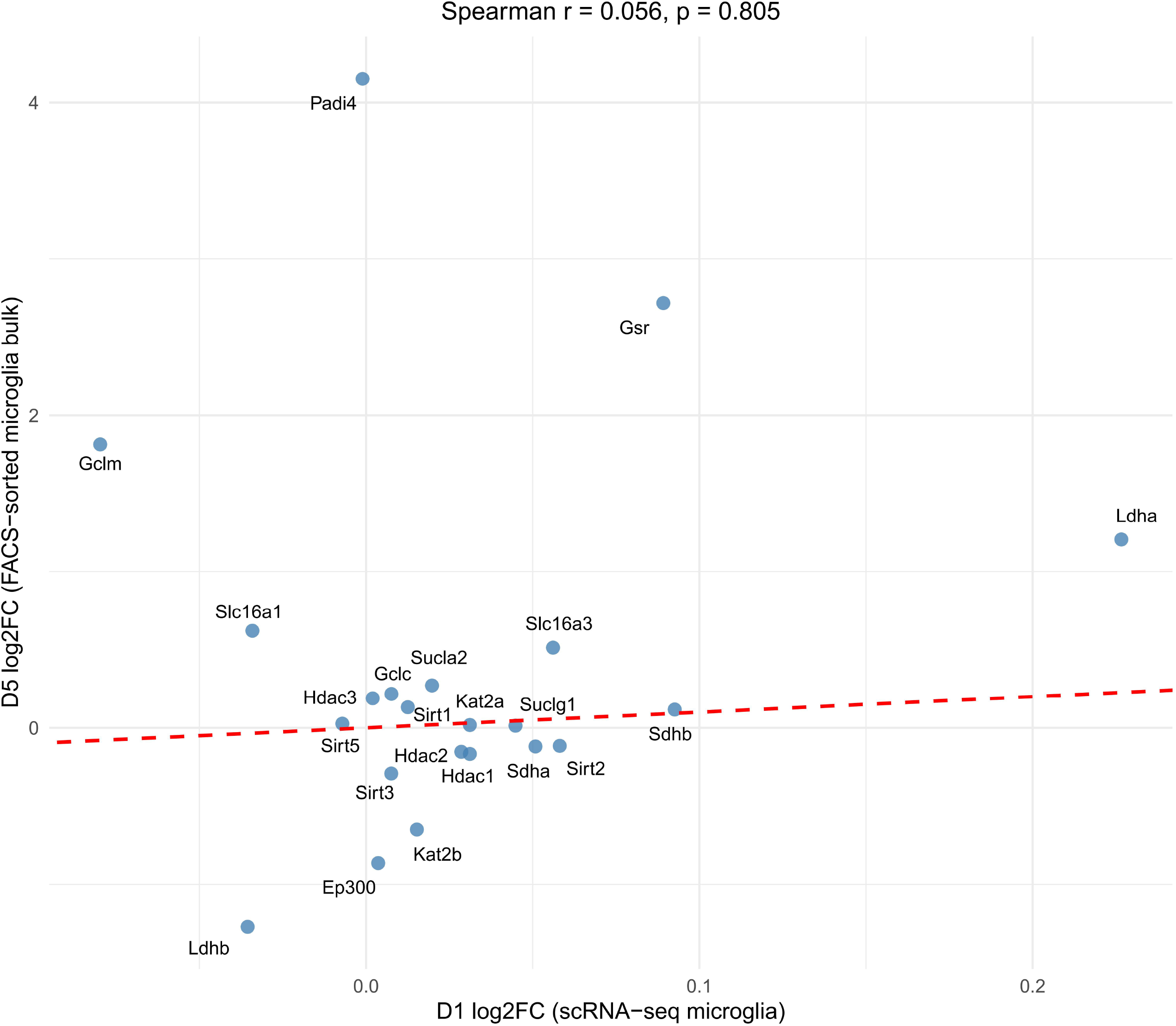
Microglial purity validation and PTM enzyme expression. (A) UMAP of D1 microglia colored by condition (Sham vs MCAO). (B) Heatmap of marker genes (P2ry12, Tmem119, Cx3cr1, Cd68, Adgre1, Ly6c). (C) Scatter plot of log2FC between D1 (scRNA-seq) and D5 (FACS-sorted bulk) for PTM enzymes (Spearman r = 0.056, p = 0.805).

## Methods

### Data acquisition and preprocessing

Raw data from GSE174574^[15]^, GSE227651^[16]^, GSE245386^[17]^, GSE267240^[18]^, and GSE319237^[19]^ were downloaded from GEO. For scRNA-seq datasets, Seurat v5.0.3^[20]^ was used for quality control, normalization, and integration. Cells were filtered for nFeature_RNA 200–6000, nCount_RNA 500–30000, percent.mt < 20%, and percent.rb < 5%. Each dataset was independently normalized using SCTransform (v2, regressing out percent.mt). Dimensionality reduction was performed with PCA (50 PCs) and UMAP (dims=1:30, seed=2026). Clustering used FindNeighbors and FindClusters (resolution=0.8).

### Microglia double filtration

Microglia were identified based on expression of P2ry12, Tmem119, and Cx3cr1 (≥2 genes positive) and absence of Cd68, Adgre1, and Ly6c (all negative) using the data layer of SCTransform. Contamination rate was calculated and verified by UMAP visualization of Cd68 expression.

### PTM enzyme gene set and NMF module identification

PTM enzymes (25 genes, Table S2) were derived from literature^[11-14, 21]^. Genes expressed in <5% of cells were filtered, leaving 22 genes. NMF^[22]^ was performed using the NMF package v0.28^[23]^ on the cell × gene matrix (SCTransform data layer). The optimal rank (k=3) was chosen based on cophenetic coefficient and biological expectation. Stability was assessed by nrun testing (20, 50, 100), 5-fold cross-validation, and sample-level validation (three individual samples). Differential expression for module marker genes was performed with FindAllMarkers (logfc.threshold=0.5, min.pct=0.25, only.pos=TRUE). GO enrichment analysis used clusterProfiler v4.10.0^[24]^ with org.Mm.eg.db^[25]^ (p.adjust < 0.05).

### Spatiotemporal analysis

The D1 weight matrix W (genes × modules) was used to project onto D2 microglia via non-negative least squares. The resulting H matrix (cells × modules) provided module scores. Module proportions per timepoint were calculated, and chi-square test assessed association. Pseudotime analysis was performed with Monocle3 v1.3.1 ^[26]^ using Sham cells as root.

### Independent validation and sex analysis

D3 (GSE245386) was projected with D1 W matrix, and module proportions were compared to D1 MCAO using cosine similarity. D4 (GSE267240) was similarly projected; sex differences in stroke group module distribution were tested with chi-square and Fisher exact test (10,000 simulations) and bootstrap resampling.

### Purity validation

D5 bulk RNA-seq data were normalized (log2CPM). PTM gene log2FC between MCAO and Sham were calculated and correlated with D1 log2FC using Spearman’s rank correlation.

### Software and reproducibility

All analyses were performed in R 4.5.2. Random seed was fixed at 2026. Session information is provided in Supplementary File 1.

## Discussion

This study provides the first single-cell transcriptomic characterization of PTM enzyme co-expression modules in microglia during CIRI. We identified three robust modules—metabolic stress, pro-inflammatory, and reparative—that exhibit distinct spatiotemporal dynamics and are validated in an independent dataset. Our findings support a “PTM enzyme code” hypothesis, where coordinated expression of PTM regulatory enzymes defines microglial functional states.

### Biological significance of modules

The metabolic stress module (M1) was enriched in TCA cycle enzymes and declined rapidly after reperfusion, consistent with a homeostatic state relying on oxidative phosphorylation^[9]^. The pro-inflammatory module (M2) peaked at 1–3 days, aligning with the acute inflammatory phase and enrichment in pathways such as leukocyte activation and chemotaxis^[3, 27]^. The reparative module (M3) showed a late slight increase, with vascular development and translation pathways, suggesting a role in tissue remodeling and recovery^[28]^.

Of note, the M3 (reparative⍰associated) module exhibited a relatively dispersed distribution in UMAP space (Fig. 3A). This likely reflects the inherently heterogeneous nature of reparative microglia, which may encompass cells at different stages of transition from pro⍰inflammatory to repair states, as well as cells with distinct functional preferences (e.g., angiogenesis vs. synaptic remodeling). The dispersed pattern, rather than a tight cluster, supports the view of microglial functional continuity and aligns with our proposed PTM enzyme code framework that moves beyond the classical M1/M2 dichotomy.

**Figure 3.**
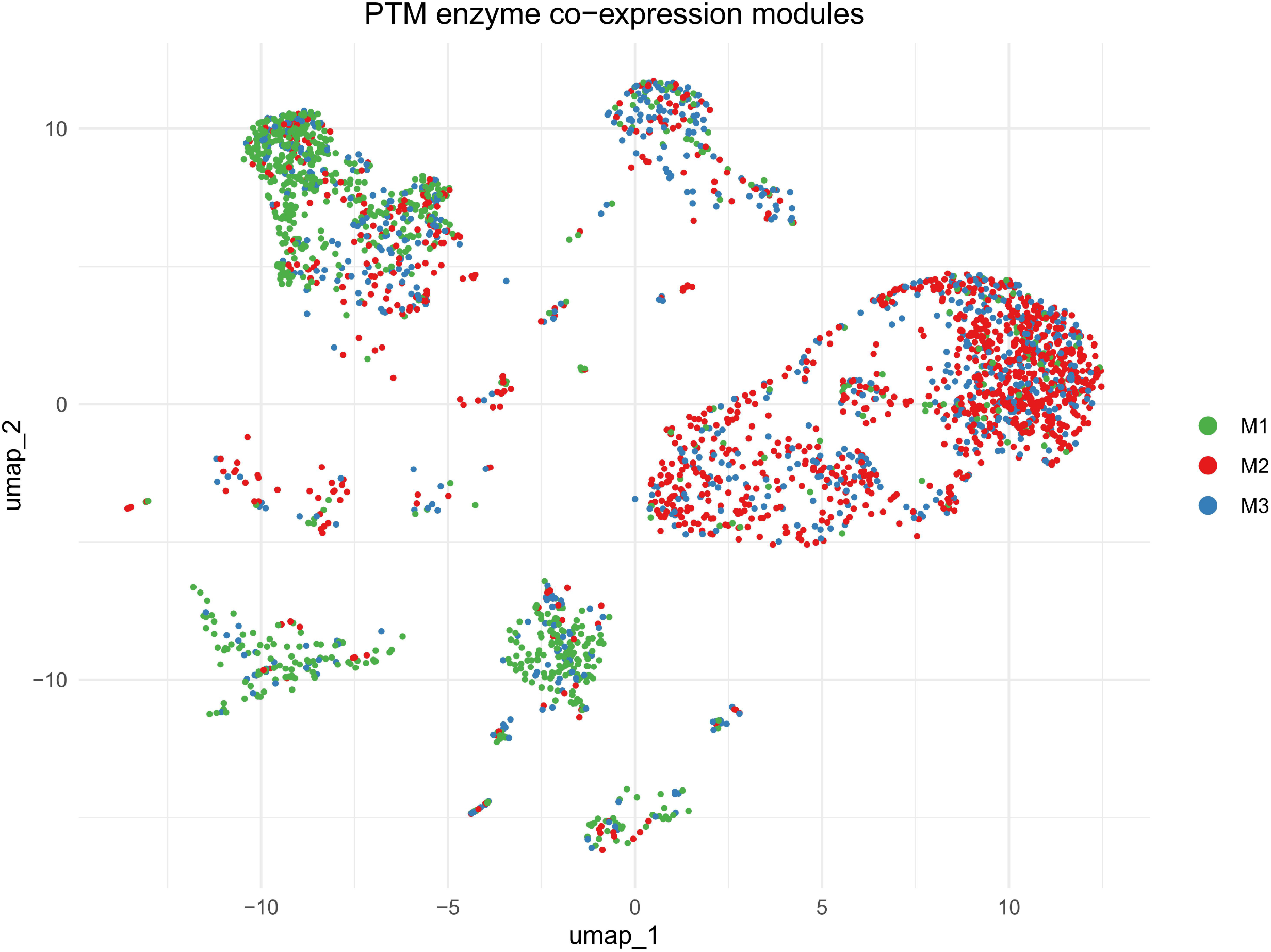
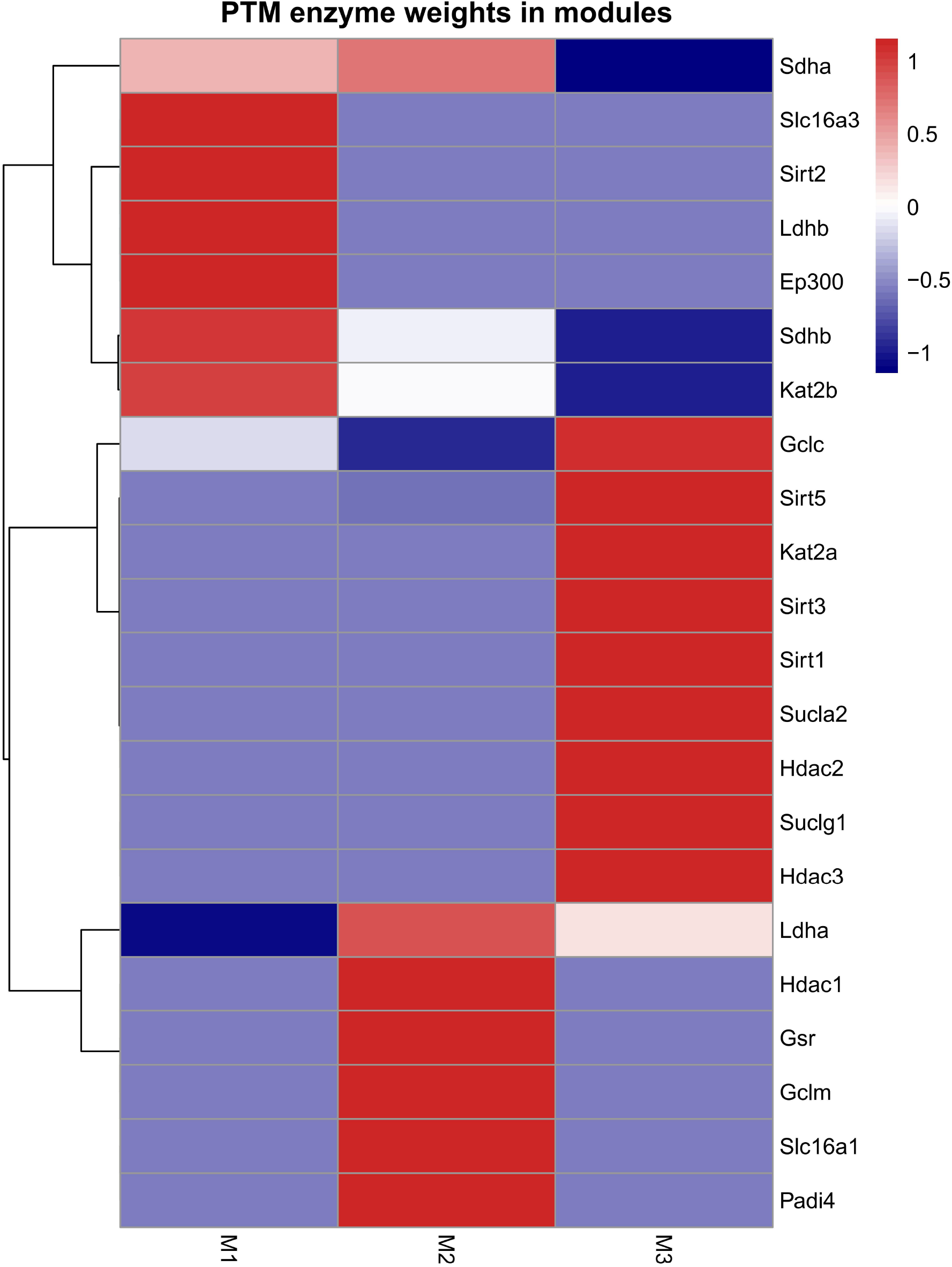
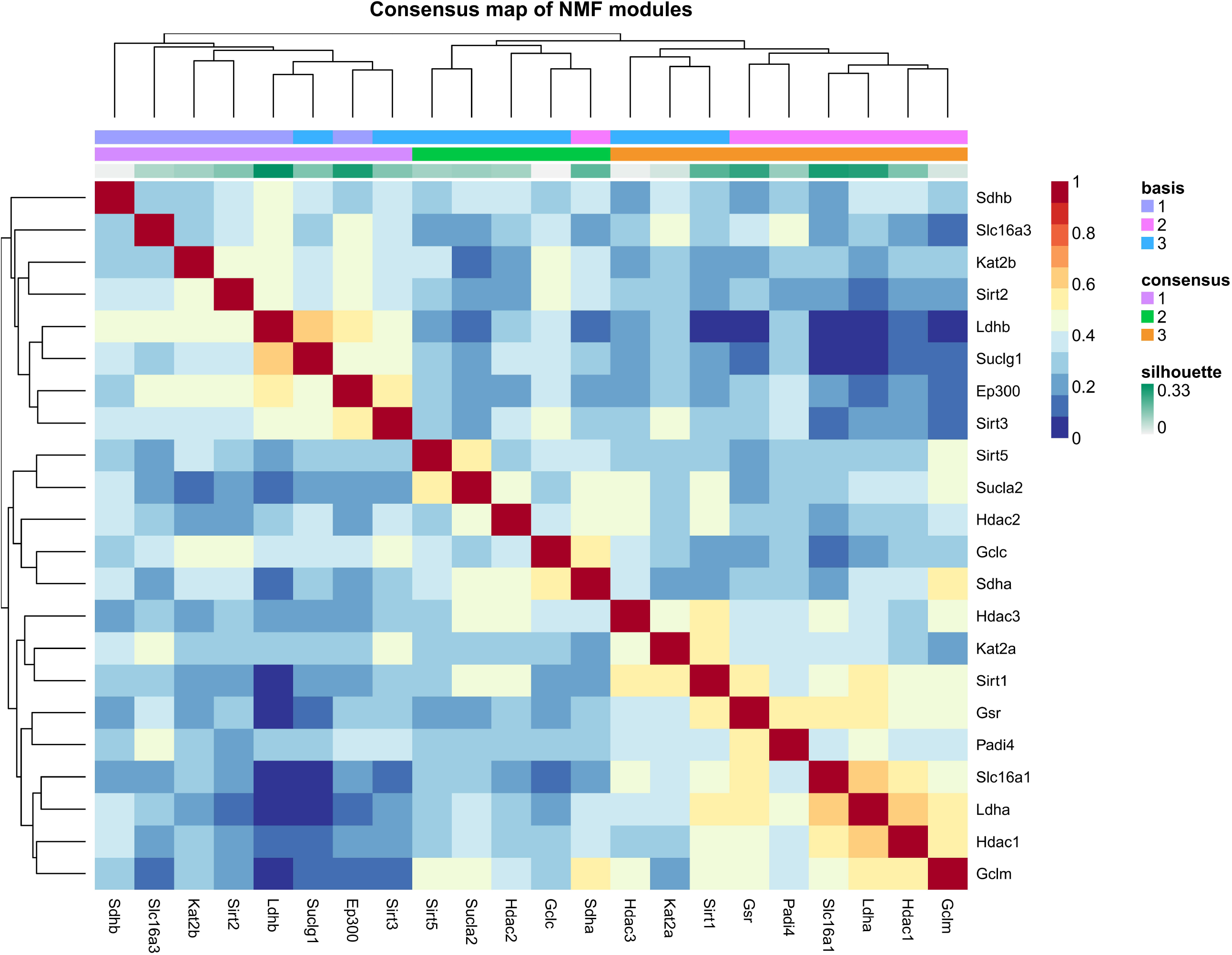
Identification of three PTM enzyme co-expression modules. (A) UMAP of D1 microglia colored by PTM enzyme co-expression module assignment (M1: metabolic stress-associated, M2: pro-inflammatory-associated, M3: reparative-associated). The M3 module shows a relatively dispersed distribution, reflecting the heterogeneous and transitional nature of reparative microglia, which is consistent with the functional continuity of microglial states beyond the classical M1/M2 paradigm. (B) Heatmap of gene weights in each module (22 genes × 3 modules). (C) Consensus map of NMF modules demonstrating stability. Sample-level validation results are described in the text.

**Figure 4.**
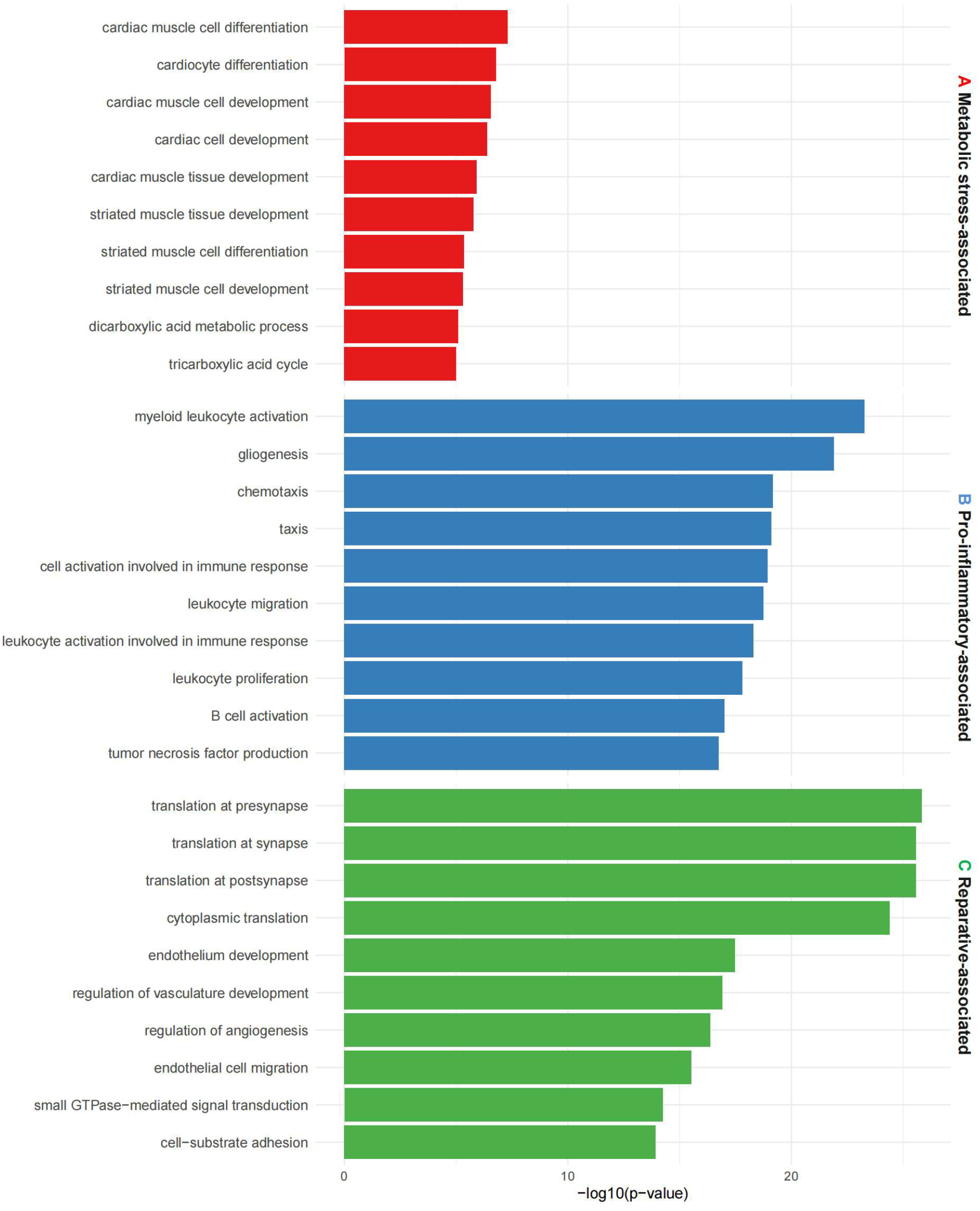
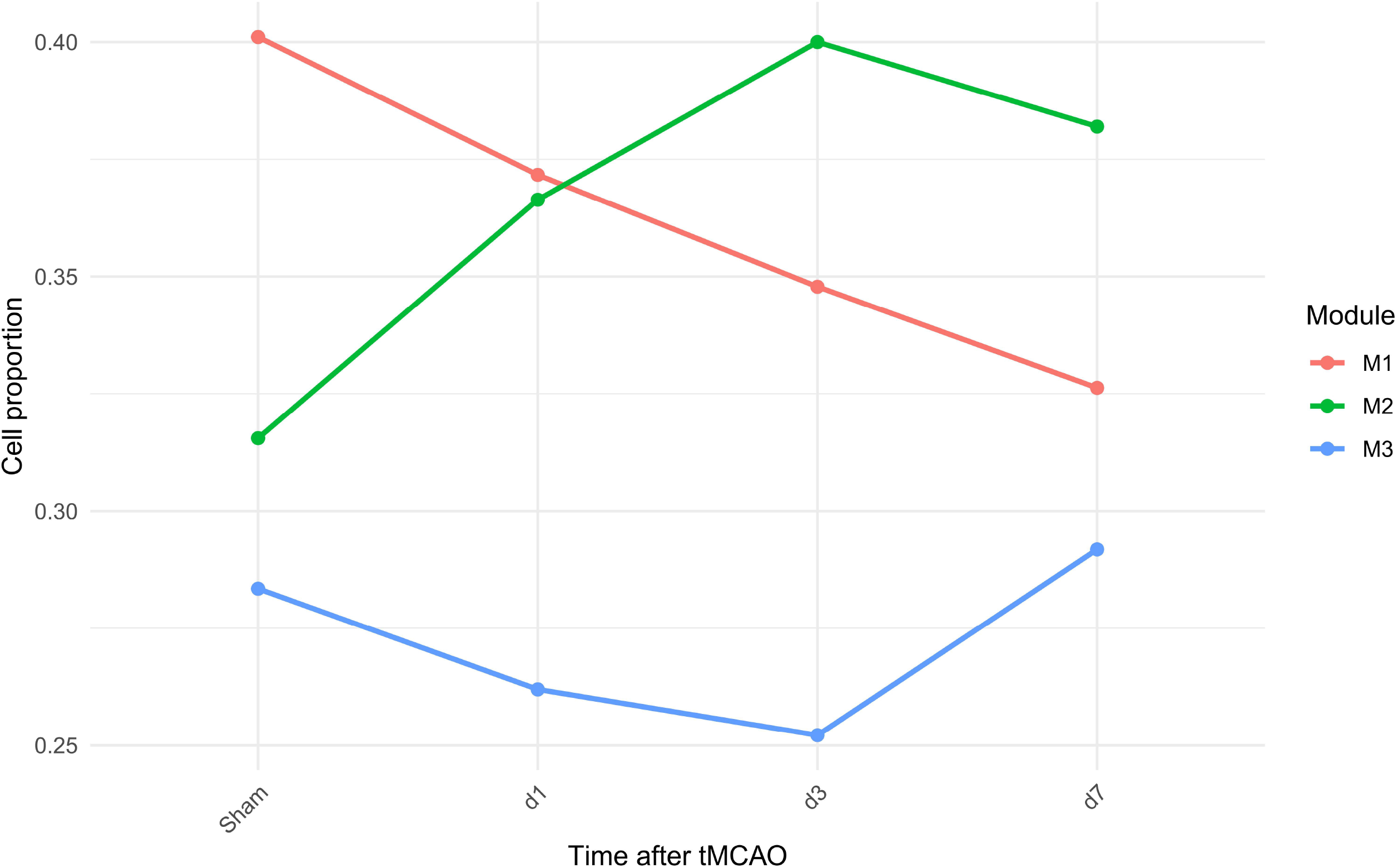
Functional annotation of modules. Top GO biological processes enriched in (A) Module 1 (metabolic stress), (B) Module 2 (pro-inflammatory), and (C) Module 3 (reparative).

**Figure 5.**
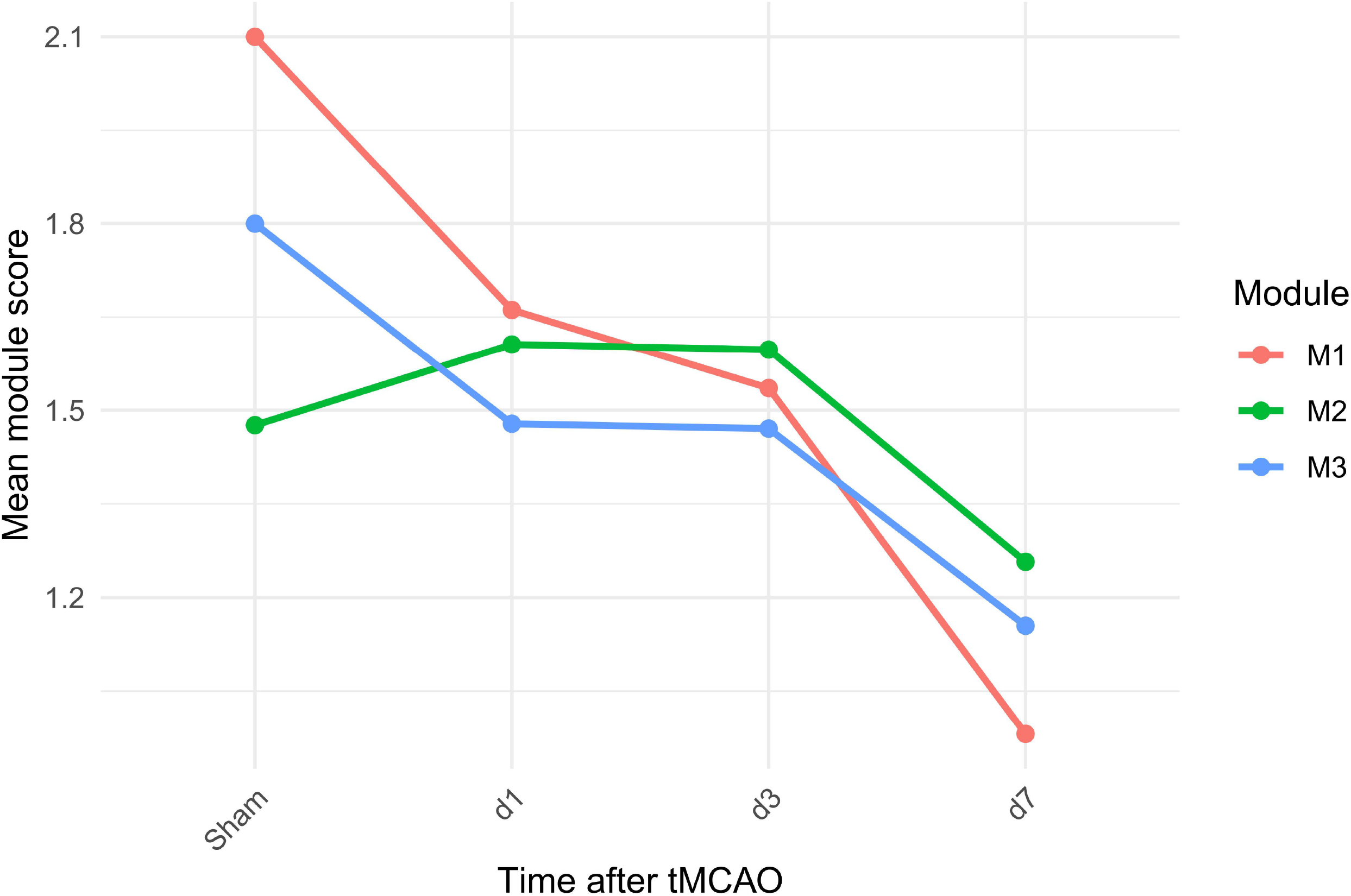
Spatiotemporal dynamics. (A) Module proportions across timepoints in D2. (B) Mean module scores across timepoints. Data are mean ± SD.

**Figure 6.**
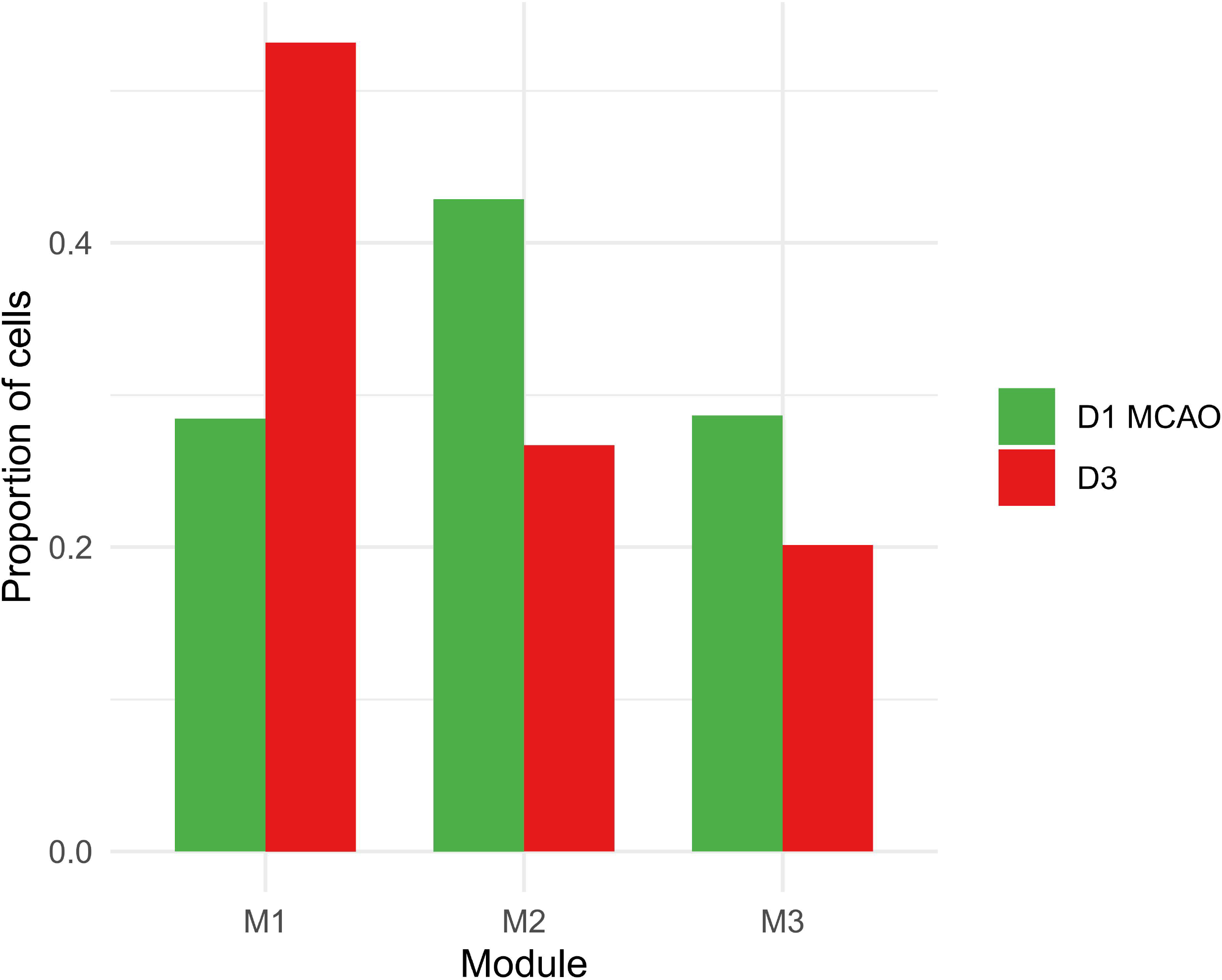
Independent validation. Module proportion comparison between D3 and D1 MCAO (cosine similarity = 0.874).

**Figure 7.**
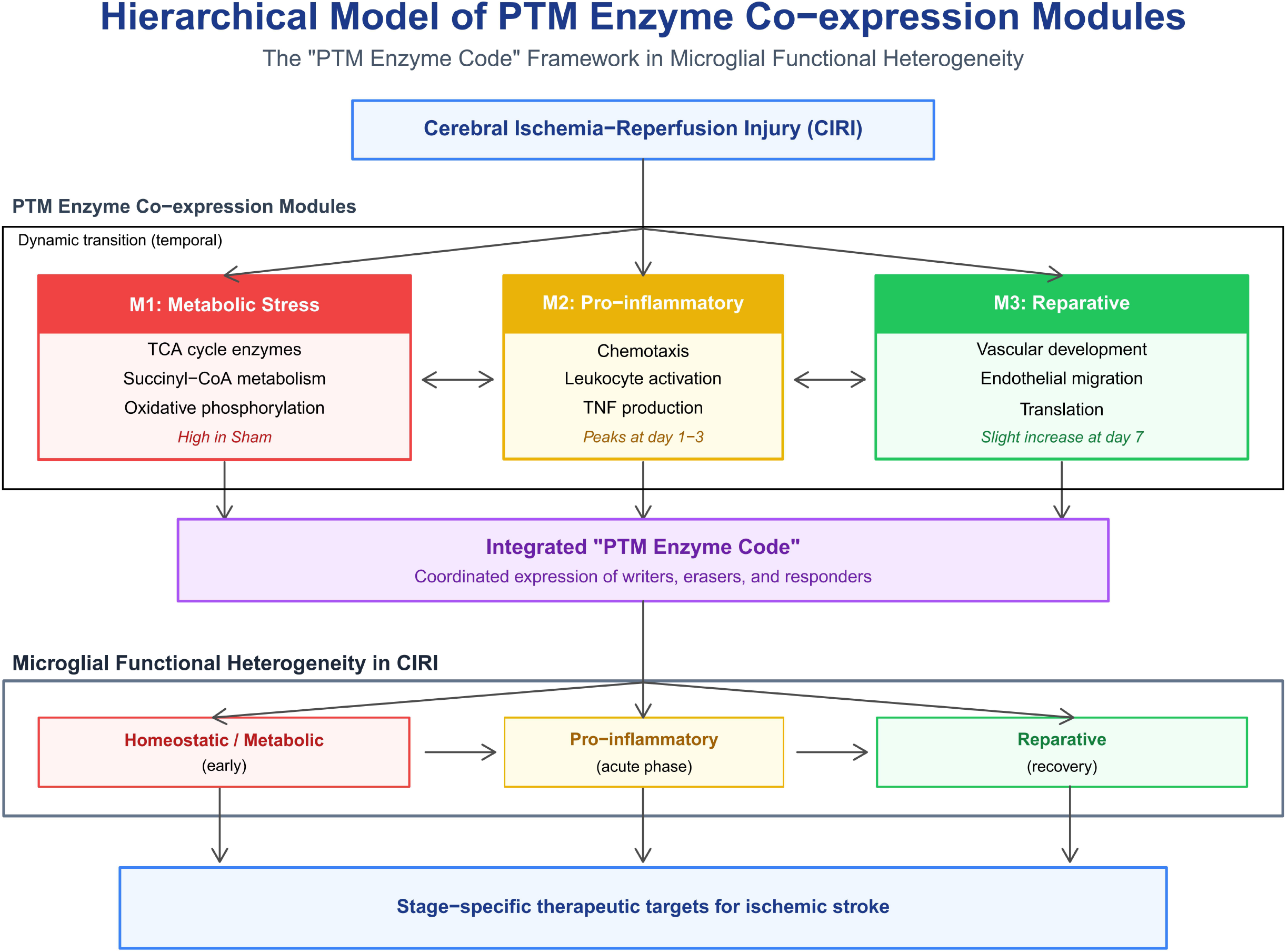
Hierarchical model of PTM enzyme co-expression modules. Schematic integrating the three modules into the “PTM enzyme code” framework.

### Spatiotemporal transitions

Despite non-significant chi-square test, the clear trends in module proportions and scores support a sequential transition from metabolic stress to pro-inflammation and later a shift toward repair. This is reminiscent of the dynamic microglial states observed in other neurological disorders^[29]^ and aligns with the proposed hierarchical “PTM code” model.

### Sex differences

The observed sex differences in module distribution in permanent ischemia are consistent with known sex-biased microglial responses^[30, 31]^, but need validation in reperfusion models. The large sample size in D4 strengthens the finding, but model differences caution against direct extrapolation.

### Purity and limitations

The low correlation between D5 bulk and D1 single-cell PTM expression likely reflects model and timepoint heterogeneity, not impurity. Our double filtration effectively removed macrophages, and UMAP showed minimal Cd68^+^ cells. Limitations include the use of enzyme expression rather than direct PTM measurement, lack of spatial resolution, and absence of D6 (due to data incompatibility). Currently, public databases lack single-cell transcriptomic data of human microglia in ischemic stroke, which limits cross-species validation of our findings. Nevertheless, we observed highly conserved PTM enzyme co-expression modules across multiple independent mouse stroke models (covering tMCAO reperfusion, pMCAO permanent ischemia, and both sexes), suggesting that PTM enzyme co-expression patterns may be universal in regulating microglial function. Future studies using human samples (e.g., post-mortem brain tissue or surgical specimens from stroke patients) are needed to further validate their clinical relevance. Future work should integrate single-cell proteomics, PTM-specific antibodies, and spatial transcriptomics to directly test the PTM code hypothesis^[32]^.

### Therapeutic implications

The stage-specific dynamics of PTM enzyme modules suggest that targeting the metabolic stress module early, the pro-inflammatory module during the acute phase, and the reparative module during recovery might be beneficial. Such approaches could leverage existing compounds (e.g., itaconate derivatives, SDH inhibitors, SIRT modulators) with improved brain delivery.

## Conclusions

PTM enzyme co-expression modules define metabolic, pro-inflammatory, and reparative microglial states in CIRI with distinct spatiotemporal dynamics. This transcriptional framework provides indirect support for the “PTM enzyme code” hypothesis and offers stage-specific therapeutic targets for stroke.

## Acknowledgements

Not applicable.

## Author contributions

Ying Li: Investigation, Data curation, Formal analysis, Writing – original draft, Writing – review & editing.

Huimin Li: Investigation, Visualization, Writing – original draft, Writing – review & editing.

Meng Zhang: Conceptualization, Methodology, Investigation, Visualization, Writing – original draft, Writing – review & editing, Funding acquisition, Supervision, Project administration.

## Competing interests

The authors declare no competing interests.

## Funding

This work was supported by the Zhejiang Province Traditional Chinese Medicine Science and Technology Project (Grant No. 2024ZF043 to M.Z.).

## Data availability

All raw data are available from GEO under accession numbers GSE174574, GSE227651, GSE245386, GSE267240, and GSE319237. Processed data and code are available at https://github.com/zh3028/CIRI_PTM_microglia.

## Notes

https://github.com/zh3028/CIRI_PTM_microglia

https://www.ncbi.nlm.nih.gov/geo/query/acc.cgi?acc=GSE174574

https://www.ncbi.nlm.nih.gov/geo/query/acc.cgi?acc=GSE227651

https://www.ncbi.nlm.nih.gov/geo/query/acc.cgi?acc=GSE245386

https://www.ncbi.nlm.nih.gov/geo/query/acc.cgi?acc=GSE267240

https://www.ncbi.nlm.nih.gov/geo/query/acc.cgi?acc=GSE319237

### Competing Interest Statement

The authors have declared no competing interest.

### Summary of Updates

Version 2 revision This revision corrects an error in the author list within the PDF. The correct first author is Ying Li. The PDF has been updated accordingly. The online metadata was already correct and remains unchanged.

