## Supplementary figures and images for "A PTM Regulatory Enzyme Co-expression Code Defines Microglial Functional Heterogeneity in Cerebral Ischemia-Reperfusion Injury"

### Figure S1A

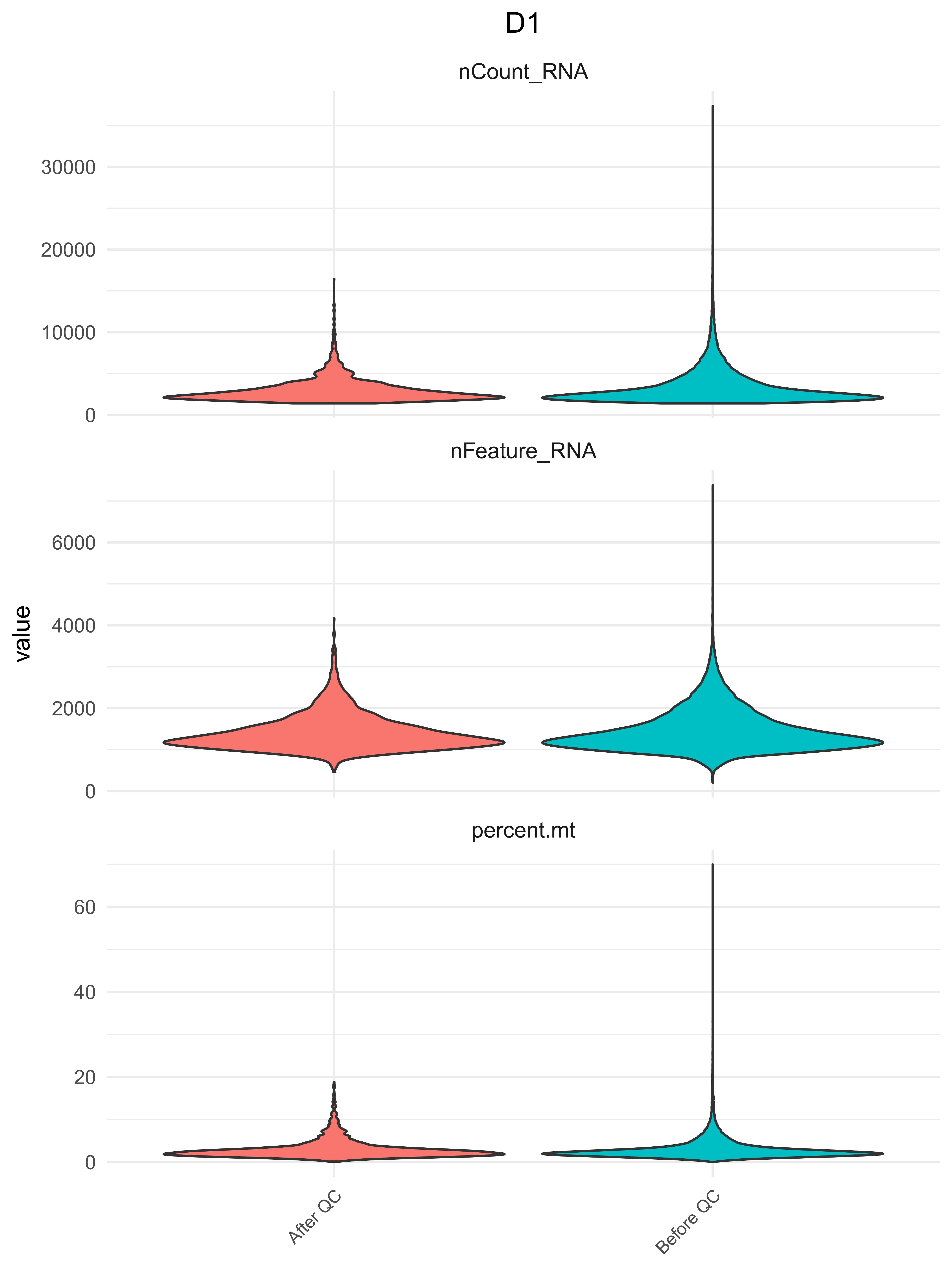

### Figure S1B

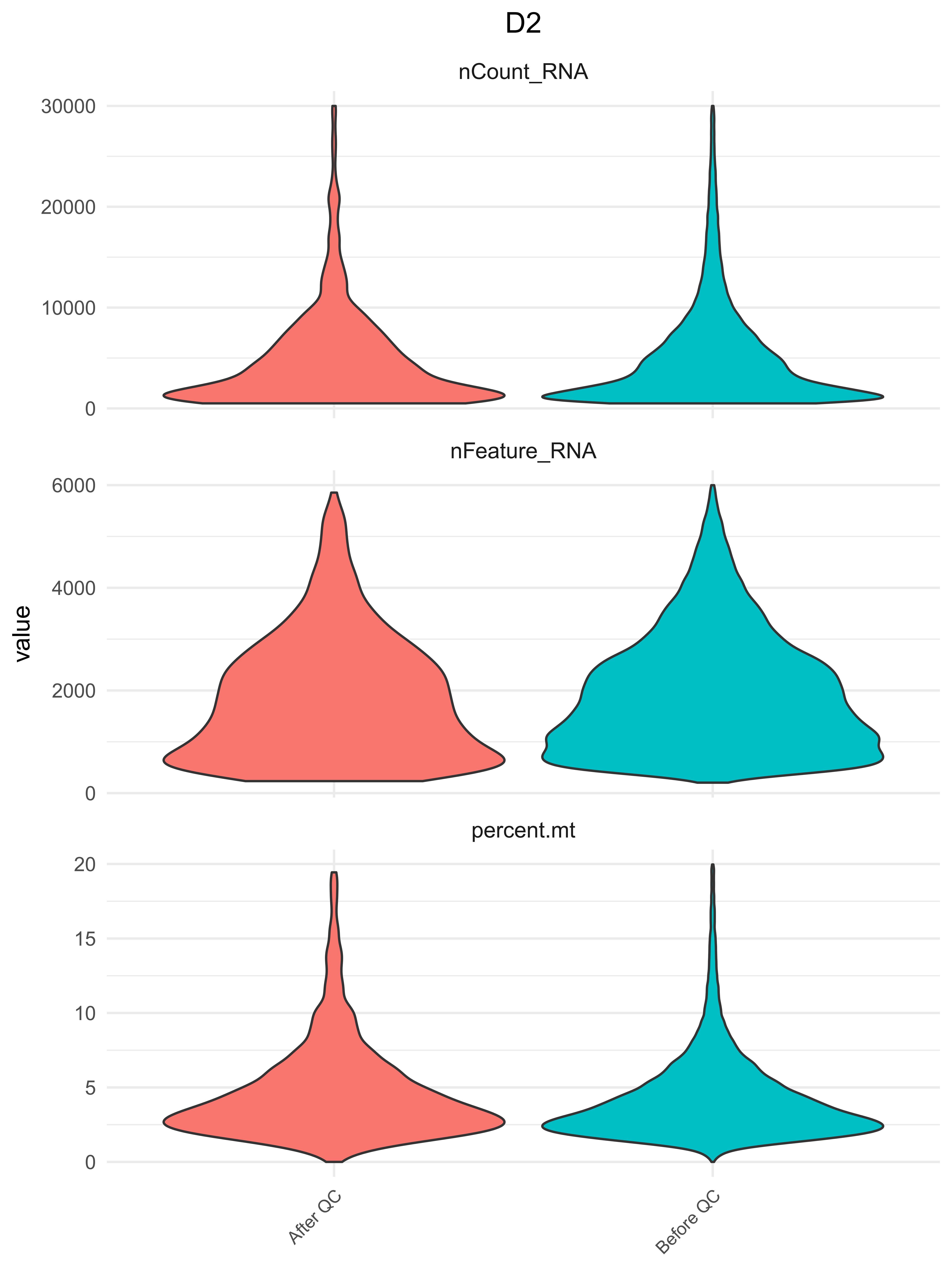

### Figure S1C

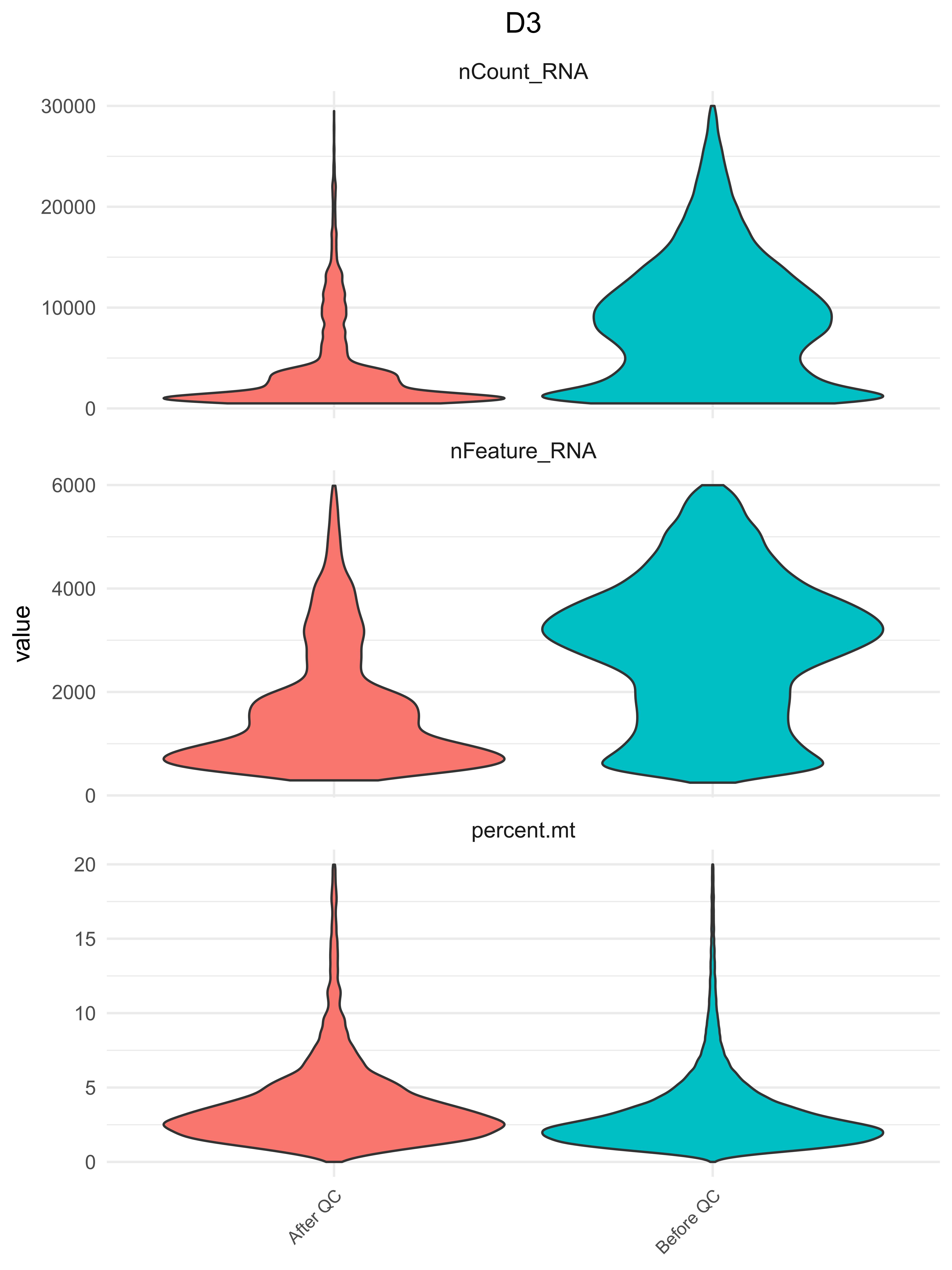

### Figure S1D

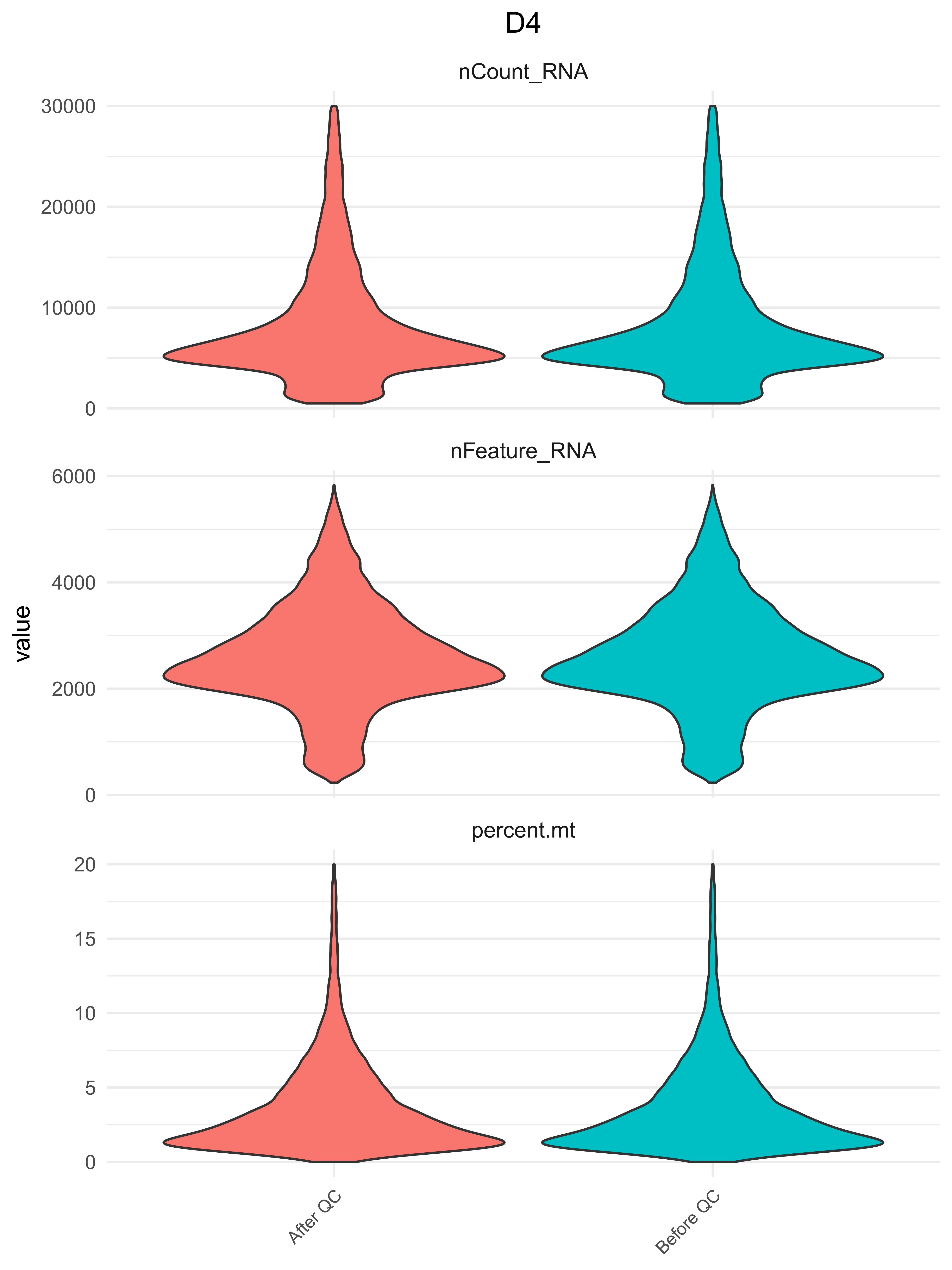

### Figure S2

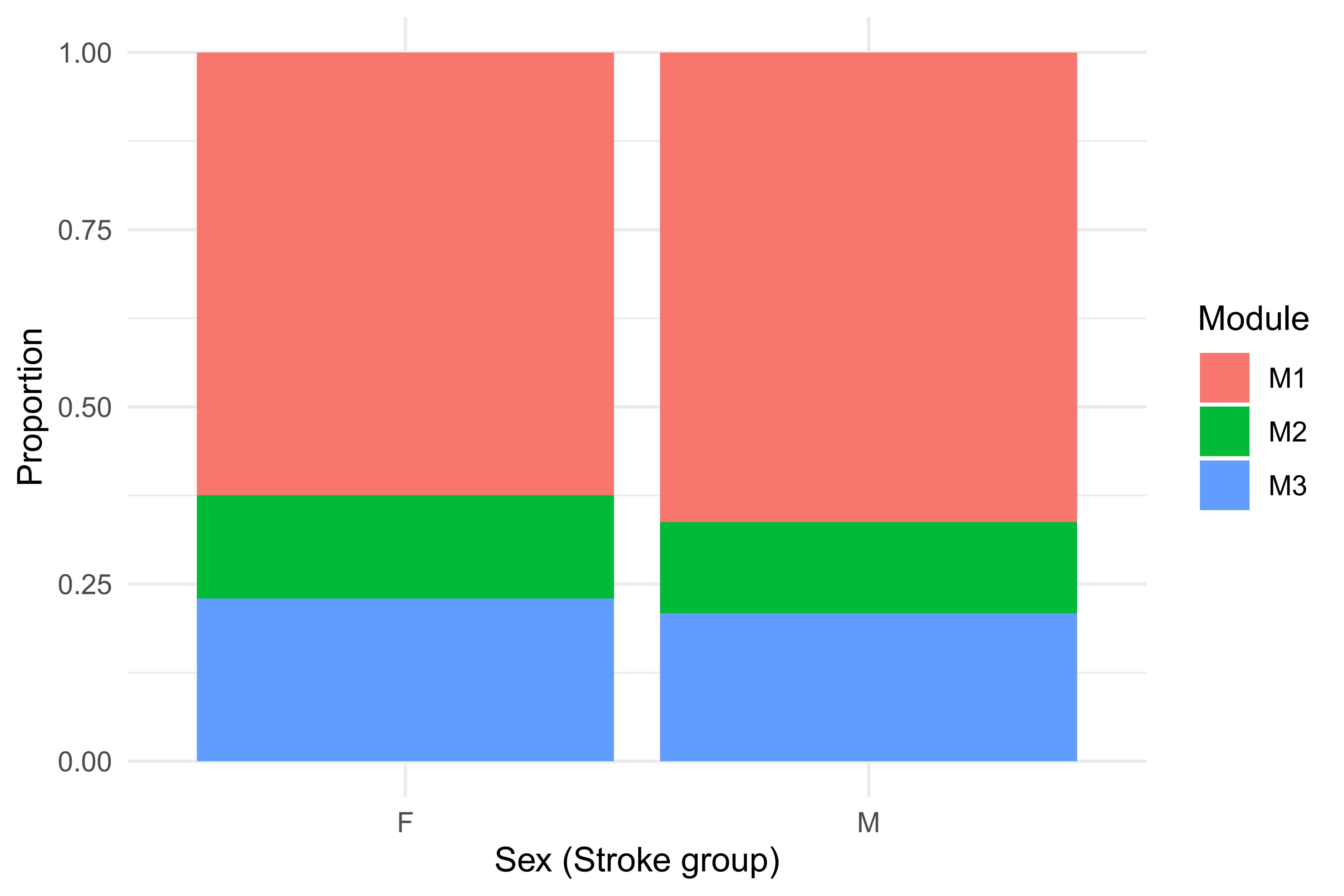
