## Supplementary material for "A PTM Regulatory Enzyme Co-expression Code Defines Microglial Functional Heterogeneity in Cerebral Ischemia-Reperfusion Injury": Table S1

| p_val | avg_log2F( | pct.1 | pct.2 | p_val_adj | cluster | gene |
|---|---|---|---|---|---|---|
| 2.60473649 | 2.5423901 | 0.42 | 0.128 | 4.8646058 | 3 | Slc16a1 |
| 1.55227238 | 1.81678705 | 0.183 | 0.075 | 2.8990239( | 3 | Adm |
| 4.4779447' | 1.67412302 | 0.101 | 0.056 | 0.08363009 | 3 | Ch25h |
| 1.1547788 | 1.56551905 | 0.141 | 0.061 | 2.1566649 | 3 | Slc19a3 |
| 1.8034037( | 1.5420666( | 0.114 | 0.048 | 3.3680368' | 3 | Unc45b |
| 1.1662440( | 1.50909748 | 0.149 | 0.062 | 2.1780773( | 3 | Rassf9 |
| 2.2906729 | 1.4580584( | 0.195 | 0.085 | 4.27806078 | 3 | Notum |
| 1.19566105 | 1.4565686' | 0.469 | 0.213 | 2.23301658 | 3 | Nostrin |
| 7.0766501 | 1.43423625 | 0.173 | 0.073 | 1.3216351' | 3 | Cdkn2b |
| 5.53073428 | 1.41916365 | 0.365 | 0.178 | 1.0329199 | 3 | Slc38a5 |
| 4.8519328 | 1.9125521 | 0.274 | 0.091 | 9.06146975 | 1 | Gsr |
| 0.0049496 | 1.8362278 | 0.1 | 0.069 | 1 | 1 | Wfdc17 |
| 2.2520587 | 1.7850394 | 0.282 | 0.092 | 4.2059449 | 1 | Hdac2 |
| 1.8544906( | 1.6534177 | 0.404 | 0.141 | 3.4634467( | 1 | Sucla2 |
| 1.9227224 | 1.50236555 | 0.461 | 0.18 | 3.59087639 | 1 | Suclg1 |
| 1.3709408 | 1.49071592 | 0.253 | 0.092 | 2.56036905 | 1 | Hdac3 |
| 0.00018429 | 1.3260374( | 0.231 | 0.172 | 1 | 1 | Gsn |
| 1.3285413 | 1.25463768 | 0.1 | 0.05 | 0.0248118 | 1 | Snhg18 |
| 1.3248600( | 1.2415465 | 0.112 | 0.051 | 0.0002474 | 1 | Anxa1 |
| 2.87115755 | 1.21634528 | 0.202 | 0.134 | 0.0536217 | 1 | Tagln |
| 9.6353492' | 2.9947251( | 0.836 | 0.199 | 1.7994978 | 2 | Ldhb |
| 5.1221592 | 2.56303889 | 0.115 | 0.022 | 9.5661446 | 2 | Slc16a3 |
| 3.74754579 | 2.3405830( | 0.146 | 0.046 | 6.9989165 | 2 | Nkain4 |
| 3.75272249 | 2.1869581( | 0.305 | 0.196 | 7.0085845 | 2 | Aldoc |
| 3.9229307 | 2.1563753( | 0.138 | 0.048 | 7.3264654 | 2 | Scg3 |
| 2.6842149( | 2.1006642 | 0.125 | 0.041 | 5.01303975 | 2 | Acsbg1 |
| 2.71699665 | 2.0909351( | 0.125 | 0.041 | 5.0742629 | 2 | Timp4 |
| 1.9579972 | 2.0647560( | 0.107 | 0.025 | 3.65675559 | 2 | Hnmt |
| 1.0218057 | 2.05688785 | 0.127 | 0.035 | 1.9083243' | 2 | Slc39a12 |
| 1.1434326' | 2.0423540 | 0.116 | 0.036 | 2.13547485 | 2 | Acsl6 |
